# Gene Transfer Agents in Plant Pathogenic Bacteria: Comparative Mobilomics, Genomic Survey and Recombinogenic Impacts

**DOI:** 10.1101/687996

**Authors:** Mustafa O. Jibrin, Gerald V. Minsavage, Erica M. Goss, Pamela D. Roberts, Jeffrey B. Jones

## Abstract

**Background:** Gene transfer agents (GTAs) are phage-like mediators of gene transfer in bacterial species. Typically, strains of a bacteria species which have GTA shows more recombination than strains without GTAs. GTA-mediated gene transfer activity has been shown for few bacteria, with *Rhodobacter capsulatus* being the prototypical GTA. GTA have not been previously studied in plant pathogenic bacteria. A recent study inferring recombination in strains of the bacterial spot xanthomonads identified a Nigerian lineage which showed unusual recombination background. We initially set out to understand genomic drivers of recombination in this genome by focusing on mobile genetic elements.

**Results:** We identified a unique cluster which was present in the Nigerian strain but absent in other sequenced strains of bacterial spot xanthomonads. The protein sequence of a gene within this cluster contained the GTA_TIM domain that is present in bacteria with GTA. We identified GTA clusters in other *Xanthomonas* species as well as species of *Agrobacterium* and *Pantoea*. Recombination analyses showed that generally, strains of *Xanthomonas* with GTA have more inferred recombination events than strains without GTA, which could lead to genome divergence.

**Conclusion:** This study identified GTA clusters in species of the plant pathogen genera *Xanthomonas, Agrobacterium* and *Pantoea* which we have named XpGTA, AgGTA and PaGTA respectively. Our recombination analyses suggest that *Xanthomonas* strains with GTA generally have more inferred recombination events than strains without GTA. The study is important in understanding the drivers of evolution of bacterial plant pathogens.

## Background

Prokaryotic mobile genetic elements (MGEs) are sources for bacterial genome evolution and adaptation (1, 2, 3, 4). This is largely because MGEs encode enzymes and other proteins for intra- and intercellular movement of genetic material, giving rise to diversity within genomes and between bacterial populations (1,2). Intracellular movement of genetic elements or more specifically transpositional recombination, often involves various forms of transposons (such as retrotransposons and DNA transposons). Intercellular movement of DNA on the other hand additionally requires the transfer of conjugative plasmids, integrated conjugative elements, bacteriophages or self-splicing molecular parasites through conjugation, transformation or transduction (1). The array of these MGEs -transposons, ICEs, conjugative plasmids, bacteriophages, self-splicing molecular parasites, etc. or different subsets thereof in a genome are often referred to as the mobilome (5,6). The mobilome has been shown to vary significantly among bacterial populations (7,). For example, whole genome metagenomic shotgun sequencing studies of the human microbiome identified significant differences in microbial mobilomes not only between distant populations in Fiji and North America but also in gene pools within Fijian sub-populations (8). Variation in specific MGEs has also been shown to drive genome evolution. An elaborate study demonstrated that expansion of insertion sequences is associated with genome reductions in species within the genus *Bordetella* (9). Similar results but on a smaller scale were also reported in the evolution of *Yersinia* species genomes (10, 11). Understanding the mobilome of microbial populations is therefore important in characterizing the evolution of new lineages or sub-populations within microbial communities.

Many described mobile genetic elements (MGEs) encode cell-to-cell transfer machinery thereby maintaining their presence, while some MGEs are defective and lack key replication ability (5). These defective MGEs however carry their own target sites for DNA binding and cleavage and are therefore important in gene transfer. An example of such a defective mobile genetic element is the gene transfer agent or GTAs that mediate recombination in bacterial genomes. GTAs are phage-like mediators of gene transfer between bacterial cells and were first reported in the early 1970s (12). It was shown at that time that strains of bacteria carrying GTAs had unusually high recombination compared to GTA-deficient strains of the same species (12). While GTAs contain genes that are related to prophages, they are different from the latter in that random packaging of the host cells’ genome does not allow for the packaging of GTA production structures, thereby limiting GTA genetic components and production in recipient cells (13).

The presence of DNA fragments which are protected in vesicles against the activity of dnase 1 and can mobilize genetic markers has been experimentally used to classify phylogenetically distinct bacteria as GTA producers (14). An annotated GTA cluster is widespread in alphaproteobacterial, but thus far only species belonging to six genera have been experimentally tested for gene transfer activity. These genera include *Rhodobacter, Brachyspira, Desulfovibrio, Bartonella, Dinoroseobacter* and the archaeon, *Methanococcus* (13,14,15). The *Rhodobacter capsulatus* GTA is the prototypical GTA, being the most characterized and has been shown to mediate horizontal gene transfer through a combination of transduction and natural transformation (14).

The genus *Xanthomonas* is a gammaproteobacteria and many species cause devastating diseases of significant crops (16,17). Four *Xanthomonas* species including *X euvesicatoria, X perforans, X vesicatoria* and *X gardneri* are known to cause bacterial spot disease on tomato and pepper (18). Strains belonging to *X perforans* have been classified as groups 1A, 1B and 2 based on phylogeny. We recently surveyed genomic recombination in an unusual lineage of the bacterial spot pathogen, *Xanthomonas perforans*, from Nigeria that showed widespread recombination in the genome (19). This new lineage represented by strain NI1 could expand these groups beyond 1A, 1B and 2 (19,20). It was initially difficult to assign the Nigerian lineage to any of these species based on multilocus sequence analyses as it showed phylogenetic signals of *X euvesicatoria* and *X. perforans* depending on the gene combination used in building the phylogeny (21). Core genome phylogeny revealed that it was more closely related to *X. perforans*, but that it shared many genomic regions with closely and distantly related *Xanthomonas* species (19). Comparative genomics identified alleles of the lipopolysaccharide biosynthetic cluster, the type three secretion system cluster, and effector genes from closely and distantly related *Xanthomonas* species (19).

In this study, we took a mobilomics approach in order to understand the genomic divergence between the Nigerian lineage and other characterized lineages of *X. perforans*, and the drivers of genomic variations in *X. perforans* in general. We focused on comparing the arrays of already characterized prophages, transposase and prophage-like elements, in the cluster of orthologous gene (COG) database. We identified lineage-specific mobilomes in *X. perforans*. Furthermore, we identified in the NI1 strain, a unique GTA cluster which bears the hallmark of a defective prophage locus. We further surveyed the genomes of plant pathogenic bacterial species for the presence of GTA. We identified in three plant pathogenic bacterial genera similar GTA clusters with unique variation. A phylogeny-based recombination detection method was used to show that lineages with GTAs could have higher inferred rates of recombination events than lineages of the same species without GTAs. Finally, attempts were made to experimentally show gene transfer using strains from the Nigerian NI1 lineage of *X perforans*. Our conclusions from this study are important in understanding a previously unobserved driver of genome evolution in plant pathogenic bacteria.

## Results

### Mobilomes in *X perforans* and Presence of Gene Transfer Agent Gene in Nigerian Lineage of *X. perforans*

From IMG Mobilome COG database consisting of 119 COGs, only 47 COGs were present in genomes in our dataset. Results are presented in Supplementary Table 1. We found patterns of conservation of some mobile elements in the different sub-divisions of *X. perforans*. The core *X. perforans* mobilome found in all strains includes genes in COG1943, COG2801, and COG2915. Phage genes belonging to COG3740 and COG4654 are only found in group 1A strains but group 1A strains lack genes belonging to COG3645, COG4584 and COG4643. No group 1B strains contain a transposase belonging to COG3316. COG3645 is only found in strains of groups 1B and 2 but the majority of group 2 strains lack COG5004 and COG5518. Strain 4P1S2, an Italian strain which formed a distinct lineage was the only strain that lacks any gene belonging to COG3497. The NI1 lineage uniquely possess COG2932, which is a phage repressor protein containing a gene transfer domain, and COG3293, a transposase.

**Table 1.** Genera of Plant Pathogens with GTA in their Genome

### COG2932 and Discovery of GTA in *X perforans* Strain NI1 Genome and Comparison with *R. capsulatus* multi-locus GTA Genome

Since COG3293 is a transposase and there are several transposases in *X. perforans* genomes, we focused our efforts on the more unique gene member of COG2932. BLAST search of the COG2932 protein in NI1, represented by the locus tag CDO09_19890, on NCBI conserved domain database showed that it contains a GTA TIM-barrel-like domain. The GTA TIM-barrel-like domain is found in the gene transfer agent protein and was initially reported to mediate an unusually high recombination in the purple nonsulphur bacterium, *Rhodobacter capsulatus* (22,23).

We named this gene *xpGTA*. A BLAST search using genes in the GTA loci of *R. capsulatus* only returned three orthologous genes in NI1 that share some level of amino acid identity at the protein level (Table 2). This included the locus CDO09_19890 (*xpGTA)* which contains the GTA domain, a hypothetical gene in *R. capsulatus* but annotated as a predicted peptidoglycan domain-containing gene in NI1 (CDO09_05760), and the phage major capsid protein, CDO09_19960 (Table 2).

**Table 2.** Comparison of XpGTA to RcGTA Genes

In addition to the NI1 *xpGTA* gene, we further describe the putative XpGTA cluster, that includes all the genes from the integrase to the small terminase found in the region where the *xpGTA* gene is found in the genome (Fig. 1). This is because the integrase is the region for phage integration while the small terminase is the region for DNA recruitment into the prohead and cutting of concatemers to generate matured viral DNA (24,25). The GTA cluster in this strain is sandwiched between a 5-methylcytosine restriction enzyme gene, and an *rha* gene upstream and a DNA adenine methylase gene downstream, genes that are suggestive of possible roles in gene regulation and lysogeny maintenance (26).

**Figure 1.** The XpGTA Cluster in NI1 showing some genes within the NI1 XpGTA cluster.

### Identification of GTA Cluster in Other Plant Pathogenic Bacteria

Analysis of other species of plant pathogenic bacteria revealed few additional strains, with a GTA cluster, out of the many published genomes. We found an *Agrobacterium* GTA, AgGTA and *Pantoea* GTA, PaGTA. GTA found in other *Xanthomonas* species retain the XpGTA nomenclature since *X. perforans* was the first xanthomonad where a GTA cluster was identified. We did not find any evidence of GTA in genomes of species of any other major plant bacterial genera such as *Dickeya, Pseudomonas, Ralstonia, Erwinia* and *Pectobacterium* (Table 1).

GTAs of both AgGTA and PaGTA loci have genes that bear the hallmark of a phage region with few regulatory genes (Figures 2A and 2B). There is variation in the organization of AgGTA and PaGTA between species and across strains. For example, a *luxR* gene is found upstream of the *paGTA* gene in *P. ananatis* but not in *P. allii* (Figure 2a). In other *Xanthomonas* species possessing GTA, the GTA loci are similar with some variation (Figure 2c). For example, the number of known and hypothetical genes between the phage tail tape measure gene and the phage terminase gene vary across species (Figure 2c).

**Figure 2.** A. The Agrobacterium GTA, AgGTA Cluster B. The Pantoea GTA PaGTA, and C. XpGTA in other *Xanthomonas* species. Cluster and Arrows with same color point to the same gene as the one described for NI1. Genes with the same color belong to the same COG.

### *gafA* is Present in AgGTA but Absent in XpGTA and PaGTA

In previous experiments, it was shown that *rcc01865* (recently renamed *gafA)* significantly regulates gene transfer activity in *R. capsulatus,* with a deletion mutant of the gene producing no detectable gene transfer (27, 28). Homologues of this gene were also found in all confirmed GTA producers, where they are flanked by a lipoyl synthase (*lipA)* upstream and GMP synthase (*gua1)* downstream (27, 28). Table 2 shows that no homologue of *gafA* (*rcc01865*) was found in NI1. However, we carried out a BLAST search and found homologues of *lipA* and *gua1* across all the genomes with XpGTA (for NI1, CDO09_21095 for *lipA* and CDO09_09580 for *gua1*) and PaGTA (for strain AJ13355, PAJ_0439 for *lipA* and PAJ_2135 for *gua1*). These genes are also found in AgGTA and we also found homologues of *gafA* in some genomes with AgGTA. For example, *Agrobacterium vitis* strain S4 has *gafA* homologue at the locus Avi_1289 (query coverage 70% and protein identity 38%, when BLAST with *gafA* of *R. capsulatus*). Unlike *R. capsulatus*, the *lipA* and *gua1* genes in *A. vitis* are however on different contigs at different loci, Avi_2123 and Avi_0327 respectively. Comparison of the *gafA* locus in *R. capsulatus* and *A. vitis* is shown in Figure 3.

**Figure 3.** Comparison of *gafA* loci in *R. capsulatus* and *A. vitis*. *gafA* is found upstream of *lipA* in *R. capsulatus* but the *lipA* is in a different locus in *A. vitis*. Arrows of the same color in each strain points to homologues of the same gene. Genes with the same color belong to the same COG.

### Average Nucleotide Identity and Inference of Homologous Recombination in GTA and Non-GTA Lineages

GTA-containing strains in *X. oryzae* pv. *oryzae* have slightly lower ANI values compared with non-GTA strains (Table 3), mirroring the pattern previously observed between NI1 and other *X. perforans* strains (19). Strain X11-5A does not have an assigned pathovar and has the least ANI when compared with either *X. oryzae* pv. *oryzae* or *X. oryzae* pv. *oryzicola*. This was not strictly the trend with *X. oryzae* pv. *oryzicola* strains, however. Two strains with GTA, CFBP7331 and CFBP7341 have lower ANI values than other strains without GTA, but CFBP2286 has a slightly elevated ANI value. Additionally, strain CFBP7342 without GTA has a lower ANI value than typical *X. oryzae* pv. *oryzicola* ANI values. In *X. vasicola* pv. *zeae,* only strain X45 has a GTA locus and it has slightly lower ANI values than other strains, mirroring NI1-*X. perforans* story and *X. oryzae* pv. *oryzae*.

**Table 3.** Average Nucleotide Identity Comparisons between GTA and Non-GTA Strains of *Xanthomonas oryzae* pv. *oryzae, X. oryzae* pv. *oryzicola* and *X. vasicola* pv. *zeae*.

Inference of homologous recombination similarly followed the same pattern as the ANI. *oryzae* pv. *oryzae* strains with GTA have more inferred recombination events in their lineage than non-GTA *X. oryzae* pv. *oryzae* strains (Fig. 4A). This pattern was not clear for *X. oryzae* pv. *oryzicola* (Fig. 3B). Clearly, the branch with CFBP7342, CFBP7341 and CFBP7331 had more evidence of inferred recombination events than other branches (Fig. 4b). Recombination inference in *X. vasicola* pv. *zeae* mirrored that of *X. oryzae* pv. *oryzae*. Only strain X45 with a GTA locus had more inferred recombination events (Fig. 4c). The reference *X. vasicola* strain NCPB2417 which is not a zeae pathovar also contains GTA locus and this may have influenced the observed recombination events.

**Figure 4.** Inference of Homologous Recombination in *Xanthomonas* species. a). The *X. oryzae* pv. *oryzae* strains with GTA, X11-5A, MA134, NAI8NI1 and MAI1 showed more inferred homologous recombination in their branches than other strains. b). The pattern is less clear in *X. oryzae* pv. *oryzicola*. Strains CFBP7341 and CFBP7331 with GTA showed elevated recombination on their branches, like CFBP7342 without GTA. c). X45 with GTA showed more inferred recombination events than all other strains without GTA. The reference *X. vasicola* strain, though not a maize pathovar, also possess GTA. Inferred recombination is shown in dark blue horizontal lines. Invariant sites are shown in light blue (the background). White lines indicate no homoplasy while range of colors from yellow to red indicates increasing degrees of homoplasy.

## Discussion

### GTA Cluster is present in Plant Pathogens in Alphaproteobacteria and Gammaproteobacteria

In genome analysis of plant pathogenic bacteria, GTA clusters were detected in a few genera. Members of *Agrobacterium* in the alphaproteobacteria possess GTA clusters which we have designated AgGTA. Sequences with *R. capsulatus* GTA-like sequences were previously identified in *Agrobacterium* species (29). Here, we show that the AgGTA loci are characterized by phage genes, as has been shown previously in other GTAs (27; Fig. 2b). Two Gammaproteobacteria genera contain species that have GTAs. We identified the *Pantoea* GTA, PaGTA, with several phage-like genes in its cluster(Fig. 2a). Curiously, there is an outer membrane protein gene in the GTA cluster which could be important in host pathogen interaction (Fig. 2a). Packaging of host-pathogen interaction regions of the genome have been previously demonstrated for the Bartonella GTA, BaGTA. (30,31)The PaGTA may be of similar function. A *luxR* gene is also present upstream of the integrase gene and may have a similar function in the regulation of GTA production as was shown recently for *D. shibae* (15). By far, most plant pathogenic species with GTA cluster are found in the Gammaproteobacteria genus, *Xanthomonas*. Strains in several species of this genus possess the XpGTA cluster or its variant (Fig. 1 and 2c). XpGTA loci architecture provides some striking observations (Fig. 1 and 2c). It is bounded by a DNA Adenine methylase gene just after the Integrase gene and a 5-methyl cytosine specific restriction enzyme just after the small terminase for most xanthomonads (Figures 1, 2c). These two genes have been shown to be involved in the regulation of lysogeny and lytic phases of several bacteriophages, in most cases preventing a lytic phase (21,26,32,33).

The absence of a majority of RcGTA genes in XpGTA may mean that the XpGTA may have evolved differently than RcGTA, the *R. capsulatus* GTA. While pseudogenized GTA sequences of other bacteria that did not match annotated ORFs in RcGTA were inferred to be suggestive of decaying RcGTA genes in previous study (29), this does not appear to be the case in XpGTA as intact integrase and small terminase are present, suggestive of a viral origin for XpGTA. This is however less clear for AgGTA and PaGTA. Combining information from recent studies in BaGTA and DsGTA, the GTAs of *Bartonella* and *Dinoroseobacter* species respectively, it might be more appropriate to think of the origin, evolution and functioning of GTA in bacterial systems to be unique in each bacterial genus or even species (15,31,). Indeed, it has been shown that RcGTA packages chromosomes randomly, BaGTA packages more host interaction genes, while DsGTA favors packaging of high GC rich regions, showing that GTA functions differently in different species and may have different evolutionary implications (15, 31). Additionally, it was proposed that a prophage without GTA may be transformed into GTA before incorporation in the bacterial genome, enabling it to transfer amplified bacterial DNA (31). We reason that the GTA in these bacterial systems may have arisen from independent acquisition of the GTA domain by prophages of these bacteria, before their integration into the bacterial system. This is also supported by the fact that two independently isolated *Xanthomonas* phages, Xp15 and Xaj5, possess genes whose protein sequences have the GTA domain (Table 1). The genes in the GTA gene cluster may therefore show little or no similarity to the genes in the RcGTA loci.

The presence of a homologue of the GTA regulator, *gafA,* in AgGTA strains is interesting and could suggest that demonstration of gene transfer activity may be more easily realizable in these strains. This is because *gafA* is a known regulator of GTA production activation and mutants of GTA producing bacterial species deficient in *gafA* lose their gene transfer abilities (27, 28). Both XpGTA and PaGTA strains do not have *gafA*.

The absence of GTA clusters in the many sequenced strains of species belonging to other plant pathogen genera such as *Dickeya, Pseudomonas, Ralstonia, Erwinia* and *Pectobacterium* suggests that GTA may be limited to certain genera of plant pathogenic bacteria or could be of recent acquisition in plant pathogens.

### Structured Mobilomes in *X. perforans*

Comparative mobilomics provided several insights into the distribution of selected MGEs in the genomes of the plant pathogen, *X. perforans*, which could potentially provide information on important drivers of genomic evolution in plant pathogenic bacteria. Some COG groups were limited to or absent from certain groups, suggesting possible roles in lineage diversification (34). For example, COG3740 is only found in *X. perforans* group 1A while COG4653 is only found in group 1A and NI1 strains. COG3740 is a phage head maturation protease (pfam04586) while COG4653 is predicted major capsid protein (pfam05065). Phage head maturation proteins are often required to produce viable phages and infectious particles (35). While our major target was for unique variation in NI1, many other phages do exist in *Xanthomonas* genomes (36,37,38,39,). All group 1A strains lack genes belonging to COG3645, COG4584, COG4643, all of which are found in prophage genomes, suggesting variation in bacteriophages found in the different groups of *X. perforans*. These COGs are however present in groups 1B and 2 strains. The unique presence of COG2932 in NI1 led to the discovery of the GTA loci in NI1, showing the power of comparative genomics in discovery of novel genomic regions of possible biological importance within a genome.

### Gene Transfer Activity in NI1

All attempts to show gene transfer activity using genetic markers in strain NI1 did not yield positive results and neither was the filtrate resistant to DNAse. The maintenance of GTA clusters in bacterial genomes for other selective benefits other than gene transfer or chromosomal recombination have recently advanced (40). It might however be that we have not optimized the conditions in which the GTA system in NI1 functions. Similar difficulties were encountered for GTAs found in some alphaproteobacteria. For example, it was shown recently that gene transfer activity could be under the control of a quorum sensing gene in *D. shibae* (15). Deletion of the CtrA-controlled *luxI_2_* gene produced strong expression of GTA in *D. shibae* than in wild-type strains (15). The sensor kinase protein CckA and minimal changes in media composition used for growing strains have also been shown to affect GTA production (41). The composition of media especially phosphate concentration as well as salinity has been shown to influence gene transfer (41,42,43). Future research should explore these and other alternatives in order to find the best way to demonstrate gene transfer activities.

### GTA and Genome Evolution: ANI and Recombination Comparisons in Xanthomonads

The ANI of NI1 has been shown previously to be in between typical ANI values of *X. euvesicatoria* and *X. perforans*, with considerably lower values than typical *X. perforans* strains when compared with the reference strain Xp91-118 (19). This study confirms similar observations in *X. oryzae* pv. *oryzae* genomes. While all *X. oryzae* pv. *oryzae* genomes without GTA were >99% identical to the reference strain PX099A, strains with GTA had >97% ANI to PX099A (Table 3). Strain X11-5A, a US strain with unassigned pathovar, was more different from the reference strain than all other strains based on ANI. This strain was isolated from fields in Texas in the United States and was shown to be weakly pathogenic and genetically distinct from other known *X. oryzae* pv. *oryzae* strains (44). However, reduced ANI values for strains with GTA were not similarly observed in *X. oryzae* pv. *oryzicola*. *X. vasicola* pv. zeae showed a similar trend as *X perforans* suggesting possession of the GTA locus may cause a genome to be more diverged from the other strains without GTA. Recombination analyses in the shared genes of *X. oryzae* pv. *oryzae* showed a similar trend as was previously inferred for NI1. Lineages with GTA have more inferred recombination events as opposed to lineages without GTA, with GTA-containing strain X11-5A showing more recombinogenic background, confirming its distinctiveness among *X. oryzae* pv. *oryzae* strains as previously reported (44). This pattern was not exactly clear in *X. oryzae* pv. *oryzicola* strains, although strains with GTA have regions of unique inferred recombination background. It may be that GTA acquisition in this system may be new to cause a distinctive divergence or other MGEs have more pronounced influence on the evolution of *X. oryzae* pv. *oryzicola*. Additional studies would be needed to understand this.

Comparing recombination in AgGTA strains was more difficult because it was difficult to come across *Agrobacterium* strains without the AgGTA cluster. For example, all *A. vitis* strains have AgGTA, and so do most *A. tumefaciens* strains. It therefore makes little sense to compare these genomes as genomes recombined through GTA activity may also lose GTA production ability (40). In that case, genomes that arose due to GTA gene transfer may not contain the GTA cluster but may, in the short term, have similar recombination background as strains with GTA clusters.

## Conclusions

Understanding the role of GTAs in the short- and long-term evolution of bacterial plant pathogens could unlock several possibilities for devising sustainable and durable management technique. If GTA is associated with high rates of homologous recombination, strains with GTA may have more gene flux or unstable genome composition in the short term. While this may initially complicate disease management, understanding this lesson could help in making better management decision, especially targeting more stable physiologically important genes. Demonstrating gene transfer and characterizing the nature of gene packaging in the laboratory could further expand the understanding and potential utilization of this phenomenon for molecular biology as a large-scale gene transfer tool. A recent study used a simulation model to show that GTA gene clusters are maintained over many million years for reasons other than gene transfer or recombination (40). Probably GTA in these plant pathogens are maintained for such additional reasons.

## Materials and Methods

### Comparative Mobilomics

Comparison of mobilomes was carried out using the cluster of orthologous genes (COG) database on the IMG platform (https://img.jgi.doe.gov/cgi-bin/mer/main.cgi). All previously published sequences of *X. perforans* genomes in the IMG database were added to the Genome cart (see supplementary text for list of *X. perforans* genomes used). COG browser was used to identify the mobilome database. The mobilome database, consisting mostly of prophages and transposons, was added to the ‘Function’ cart. Function comparison of all the genomes against genes in the Function cart was carried out by BLAST. This was achieved by using the ‘Function Comparison’ under the ‘Abundance Profile’ in the Compare Genomes tab. The tabulated output was confirmed manually and variation in COGs present in each strain was identified. COG assignments unique to the Nigerian lineage were further investigated.

### Identification of Gene Transfer Agent in Nigerian Strain of *X. perforans*

We explored the gene identified as a member of COG2932 which was unique to NI1 by carrying out BLAST on the NCBI conserved domain database. The protein sequence of the gene contained a gene transfer agent domain. We further explored this gene and the cluster of genes up and downstream of the GTA cluster of NI1, delimiting it with phage-related genes as found in the prototypical GTA bacterium, *R. capsulatus* and other phages. We further compared GTA genes in the prototypical GTA, *R. capsulatus*.

### Identification of GTA Loci in other Plant Pathogen Genomes

We further searched several genome databases of major plant pathogenic bacteria for the presence of GTA domain containing gene. The databases include IMG, EDGA (https://edgar.computational.bio.uni-giessen.de), NCBI whole genome contigs BLAST, *Pseudomonas syringae* database (http://www.pseudomonas-syringae.org/). The major genera for which search was conducted include *Agrobacterium, Pseudomonas, Xanthomonas, Ralstonia, Erwinia, Dickeya, Pantoea* and *Pectobacterium*. We compared the gene neighborhood organization of the GTA loci in strains with GTA.

### Average Nucleotide Identity (ANI) and Inference of Genomic Recombination in GTA and non-GTA strains of the same Species

We have shown previously that strain NI1 has an unusual ANI value and higher inferred recombination events compared to other *X. perforans* strains. We used publicly available genomes of strains of plant pathogenic bacteria in the NCBI database with and without GTA to test this pattern of ANI variation and infer the potential differences in genomic recombination between the two sets of strains of a species. We found several species with sequenced genomes that could be sampled for such studies. Inference of ANI and genomic recombination was therefore limited to *X. oryzae* pv. *oryzae*, causal agent of bacterial blight of rice; *X. oryzae* pv. *oryzicola*, causal agent of bacterial leaf streak of rice and *X. vasicola* pv. *zeae*, causal agent of bacterial leaf streak of maize. We included the reference strain for each species to provide evolutionary context.

The core genes of each set of strains were extracted and aligned using the Roary pipeline which first aligns individual core genes before concatenation (45). Concatenated alignment was used as the input file for iQtree (46) to infer maximum likelihood phylogeny. To infer recombination in the core genome using ClonalFrameML (47), the transition/transversion ratio for each set of alignment, a midpoint rooted Newick tree, and core genome alignment were fed into ClonalFrameML to infer recombination.

## Declarations

## Acknowledgements

The authors are grateful to the editor and reviewers for constructive comments that helped improve the manuscript.

## Authors’ Contributions

MOJ, JBJ, PDR and EMG conceived and coordinated the project. MOJ, GVM, JBJ carried out laboratory experiments. MOJ and JBJ carried out bioinformatics research. MOJ wrote the paper with contributions from JBJ, PDR and EMG. All authors read and approved the final manuscript.

## Competing Interests

The authors declare that they have no competing interests.

